# A Self-Priming Neural Chain Links Sequential Behaviors Across Timescales

**DOI:** 10.64898/2026.06.02.729657

**Authors:** Evan S. Hill, Ion R. Popescu, Jean Wang, William N. Frost

## Abstract

Behavioral sequences are essential for survival, yet the neural mechanisms that link one action to the next remain incompletely understood. In classical chain models, sequential behaviors arise through feedforward propagation of activity across distinct neuronal populations or network modules. Here, we identify a distinct form of neural chain mechanism in which neurons active during a first behavior modulate themselves into a persistent state of elevated tonic firing that subsequently drives the second behavior from within the first circuit module. We term this process self-priming, reflecting the role of activity-dependent auto-modulation in enabling behavioral sequencing.

We investigated this mechanism in the escape swim–crawl sequence of the marine mollusk *Tritonia diomedea*, in which rhythmic swimming is consistently followed by tens of minutes of rapid crawling. Serotonergic dorsal swim interneurons (DSIs), components of the swim central pattern generator, were previously known to exhibit elevated firing for tens of minutes after a swim motor program (SMP) and to drive crawling. We show here that their sustained post-SMP activity arises from self-induced increases in DSI excitability: their bursting during the SMP produces long-lasting depolarization and enhanced excitability within the DSI population. This persistent state drives prolonged elevated tonic firing which, in turn, drives escape crawling.

Our findings demonstrate a self-priming neural chain mechanism in which activity during one behavior generates the internal drive for the next. Unlike previously described feedforward chain mechanisms, sequencing in this system does not depend on sequential recruitment of distinct neural substrates. Instead, the same neurons participate continuously across both behaviors while their activity-dependent change in state links the two actions into a coordinated sequence. This mechanism provides an elegant and generalizable solution for linking sequential actions across timescales.

## INTRODUCTION

Behaviors are often executed in precise, stereotyped sequences. How such sequences, many of them critical for survival, are generated by the nervous system remains a fundamental question. Sherrington and others proposed that behavioral sequences might arise from chains of reflexes in which the sensory consequences of one action trigger the next, producing a self-generating feed-forward cascade (Clower 1998; Sherrington 1947). The later discovery of central pattern generators (CPGs)— neural networks capable of generating extended behavioral programs without sensory feedback— shifted attention toward entirely central mechanisms capable of producing sequential neural activity (Delcomyn 1980). Yet despite decades of work on motor circuits, the cellular mechanisms that link one motor program to the next remained poorly understood.

Several different mechanisms for generating behavioral sequences have received experimental support in recent years (see Discussion). In one class of models, known as neural chain mechanisms, sequential behaviors arise through feedforward propagation of activity across distinct neural populations or network modules, each contributing to one component of the sequence while activating the next. Examples of such feedforward neural chain sequences have been identified in several species. In the zebra finch forebrain, sparse sequential activation of neurons within the premotor nucleus HVC has been proposed to generate birdsong syllables through a synaptically connected feedforward chain (Long et al. 2010), while heterogeneous local axonal conduction delays help shape the temporal structure of the sequence (Egger et al. 2020). In larval zebrafish, a startle response consisting of a rapid body turn followed by escape swimming is generated though sequential activation of three distinct brain regions (Xu et al. 2021). In larval *Drosophila*, crawling behavior is generated by intersegmental propagation of activity through chains of excitatory and inhibitory interneurons that coordinate sequential body contractions (Fushiki et al. 2016). Collectively, these studies support the idea that behavioral sequences can emerge through propagation of activity across sequentially recruited neural substrates, with each module triggering but otherwise not actively maintaining the next module in the sequence.

The marine mollusk *Tritonia diomedea* provides an attractive model for investigating mechanisms of behavioral sequencing because its central nervous system contains relatively few neurons, many of which are individually identifiable across preparations. These neurons generate several simple behaviors—including feeding, escape swimming and crawling—whose neural substrates can be examined in reduced preparations (Audesirk 1978; Getting 1983b; Popescu and Frost 2002; Willows 1980). Following an aversive stimulus to the tail, *Tritonia* produces a rhythmic escape swim that is followed by several minutes of rapid escape crawling (**Figure 1A**) (Audesirk 1978; Popescu and Frost 2002). While the cellular mechanisms underlying the swim and crawl motor programs are well characterized, the mechanism linking these two behaviors into a coordinated sequence has remained unknown.

**Figure 1.**
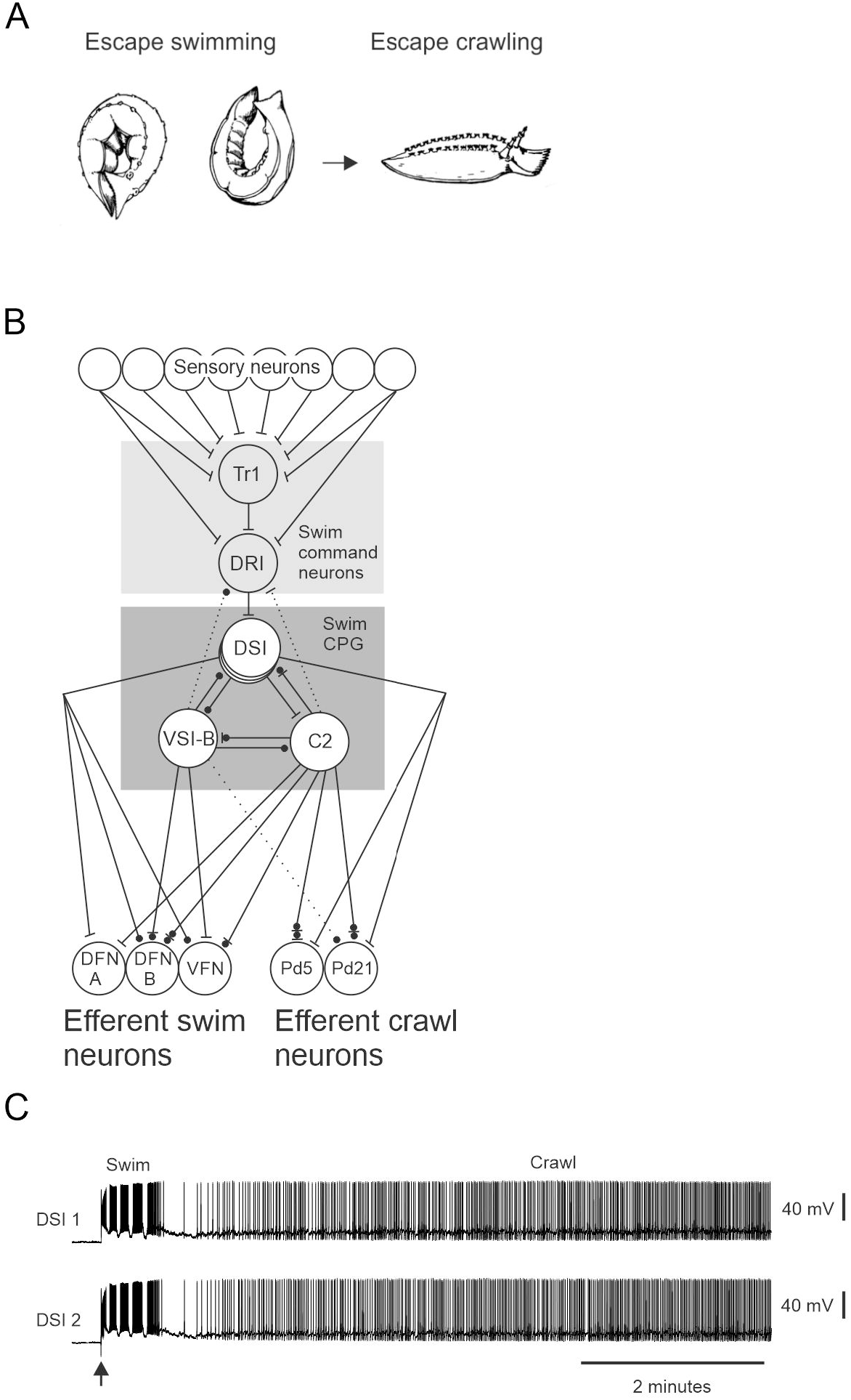
The *Tritonia* escape swim is followed by escape crawling, and the neural circuit underlying each is well-understood with one member of the swim network, the DSIs, firing rhythmically during the swim and then tonically during the escape crawl. **A** An attack by a predatory sea star triggers *Tritonia* to launch an escape swim that is always followed by tens of minutes of escape crawling. **B** Diagram of the escape swim and crawl circuits. The swim CPG consists of three types of neurons: the DSIs, C2 and VSI-B. The swim CPG neurons excite and inhibit the swim flexion neurons (the DFN-As, DFN-Bs and the VFNs). Pedal neurons 5 and 21 (Pd 5 and Pd 21) are known *Tritonia* crawling neurons. The DSIs excite both Pd 5 and 21, and C2 has complex synaptic interactions with both. Bars represent excitatory connections, circles denote inhibitory connections, and dashed lines represent polysynaptic pathways. **C** Following the rhythmic swim motor program (SMP) the DSIs fire tonically at an elevated rate for tens of minutes. The arrow denotes the stimulus to PdN 3 (10V, 10 Hz, 2 s).

The two behaviors that comprise this sequence are strikingly different (Popescu and Frost 2002). Escape swimming consists of several cycles (typically ∼5 to 8) of alternating whole-body dorsal–ventral flexions generated by the swim CPG. In contrast, escape crawling is a non-rhythmic gliding behavior produced by the beating of foot cilia and typically persists for tens of minutes (Audesirk 1978). The elements of the swim CPG were established many years ago (**Figure 1B**) (Getting 1983b). More recently, it was discovered that two components of this circuit also contribute to the subsequent crawling behavior. The neuron C2 drives the initial phase of crawling (∼30 s) (Hill et al. 2024), after which the serotonergic dorsal swim interneurons (DSIs) sustain escape crawling through prolonged tonic firing that excites crawling motor neurons (**Figure 1B–C**) (Popescu and Frost 2002). However, the mechanism responsible for generating the prolonged post-swim motor program (SMP) DSI firing, and thus linking the two behaviors into a sequence, remained unclear.

Here we show that bursting of the DSIs during the SMP induces a long-lasting depolarization in the DSIs themselves that increases their excitability and drives their prolonged tonic firing after the swim, thereby generating escape crawling. These findings identify a distinct form of neural chain mechanism in which sequential behaviors arise not through feedforward recruitment of successive neuronal populations, but through persistence-dependent transformation within a continuously active shared modulatory population. Activity during the first motor program effectively “primes the pump” for generation of the next behavior in the sequence.

## METHODS

### Animals

*Tritonia diomedea* were obtained by scuba divers off the coast of British Columbia, Canada by Living Elements. *Tritonia* were kept in tanks in our laboratory containing recirculating artificial seawater (Instant Ocean) maintained at 9.5 to 11.0° C.

### Solutions

Intracellular recording experiments were mostly carried out in Instant Ocean filtered artificial seawater (fASW). To minimize polysynaptic connections, in some experiments we used a high divalent cation saline solution (Hi Di) which had the following composition: NaCl (285 mM), KCl (10 mM), CaCl2 (25 mM), MgCl2 (125 mM), sucrose (11mM), and Hepes (10 mM); or NaCl (288 mM), KCl (10 mM), CaCl2 (29 mM), MgCl2 (129 mM) and Hepes (10 mM). Hi Di was buffered to pH 7.6 with 0.1M NaOH.

### Dissection

Dissection of the *Tritonia* central ganglia was performed in fASW, at about 7 or 8° C, and the ganglia were then dipped into a solution of 0.5% glutaraldehyde in fASW for 5 to 10 s prior to desheathing to minimize movement of the preparation during extended intracellular recordings. Desheathing was performed with fine forceps and scissors (Fine Science Tools) at 4 to 5° C.

### Intracellular recording

Borosilicate glass microelectrodes pulled by a Sutter P-87 puller were filled with 3M KCl and had resistances of 10 to 20 MΩ. DSIs were identified by location, size, appearance, spike shape and their activity during the SMP. Neuronal penetration was achieved using Sensapex uMp-3 micromanipulators. Electrodes were connected to Dagan IX2-700 dual intracellular amplifiers, and the resulting signals were digitized at 2 kHz with a BioPac MP 150 data acquisition system and recorded in the associated AcqKnowledge software, version 3.9.1. For tests of DSI excitability, 0.8 nA 5 s depolarizing pulses were injected into cells, three times roughly 5 s apart. For driving DSIs at particular frequencies for particular amounts of time (e.g. 20 Hz for 10 to 60 s) 2 ms, ∼20 nA pulses were injected into DSIs and elicited action potentials were observed on an oscilloscope to ensure that they fired spikes on all current pulses. Recordings were performed with the preparation between 9.5 and 10.5°C. Nerve stimuli were achieved via a suction electrode placed on pedal nerve 3 (PdN 3). In order to elicit an SMP, PdN3 was stimulated by a 10 V, 10 Hz, 2 s train.

### Statistical analyses

All statistical analyses were performed in Sigmaplot 11, including ANOVAs, and unpaired t-tests.

## RESULTS

### Cellular basis of the chain-sequencing mechanism

Previous treadmill experiments with intact animals have shown that *Tritonia* escape crawling can be driven by direct stimulation of the CPG’s DSI neurons (Popescu and Frost 2002), which themselves make monosynaptic excitatory connections to identified pedal crawling efferent neurons Pd 5 and 21. The DSIs burst during the SMP, during which they activate swim flexion neurons, and then the DSIs fire tonically at an elevated rate post-SMP for tens of minutes to mediate escape crawling (Popescu and Frost 2002). These observations are consistent with either 1) a chain sequencing mechanism, where the bursting of the DSIs during the SMP itself induces their prolonged elevated tonic firing to drive post-swim escape crawling, or 2) a parallel activation mechanism, in which a second, sensory-activated pathway, uninvolved in CPG oscillation, acts to induce the elevated DSI tonic firing that drives post-swim escape crawling.

To distinguish between these possibilities we tested whether bypassing the sensory neurons and directly driving the swim command neuron DRI (dorsal ramp interneuron), which strongly and monosynaptically excites all six DSIs to elicit the SMP (Frost and Katz 1996), could also elicit the elevated post-SMP DSI firing. SMPs elicited by driving DRI at 10 or 20 Hz for 40 s were followed by a significantly enhanced DSI tonic firing rate, lasting at least 5 minutes (one way repeated measures ANOVA, F (5, 3) = 9.63, p < 0.001, with post-hoc pairwise comparisons (Holm-Sidak method) over the pre-DRI spike train firing rate, N = 4 preparations); **Figure 2A, B**). This finding that bypassing the sensory neurons by directly eliciting the swim rhythm with the swim command neuron DRI induces long-lasting post-SMP elevated tonic firing in the DSIs is consistent with the CPG itself triggering the second behavior in the sequence – making this a chain sequencing mechanism.

**Figure 2.**
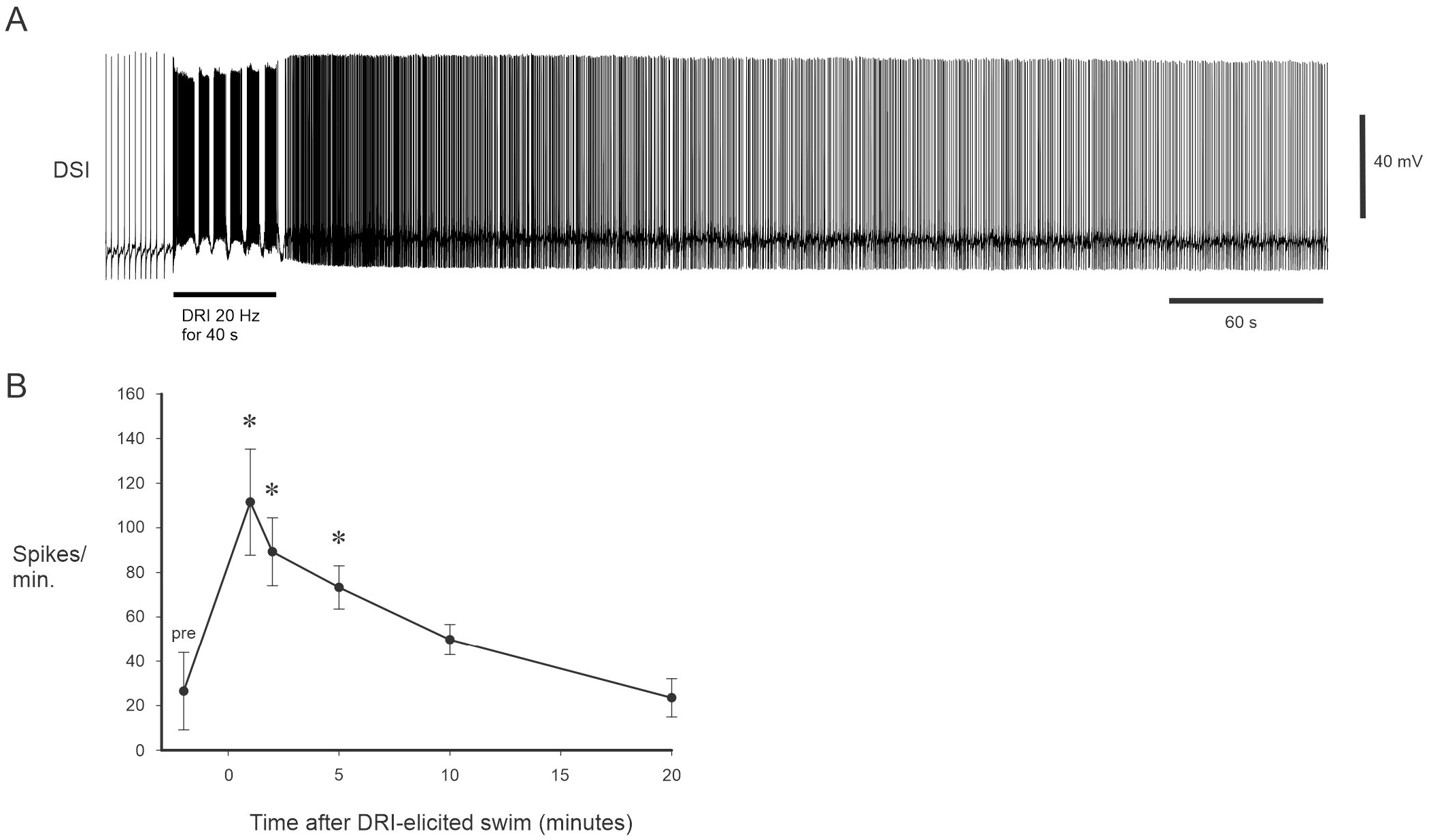
The DSIs fire at an elevated rate following SMPs elicited by driving the swim command neuron DRI, in the absence of any sensory input. This is strong evidence against parallel swim crawl circuits, and supports the notion that the escape swim and crawl behaviors are generated by a neural chain. **A** Driving DRI at 20 Hz for 40 s elicited an SMP, which was followed by elevated tonic firing of a DSI for many minutes. **B** The DSIs fire at a significantly elevated rate for 5 minutes following an SMP elicited by a DRI spike train. These data show that sensory input is not necessary for the elevated post-SMP DSI firing to occur.

We next addressed which CPG member is responsible for inducing the prolonged post-SMP firing in the DSIs: C2, VSI-B, or the DSIs themselves? Of these possibilities, the serotonergic DSIs were the most likely candidates to play this role, since they monosynaptically excite each other (Getting 1981), whereas VSI-B inhibits them (Getting 1983a), and C2’s monosynaptic connections to the DSIs are mixed but are largely inhibitory (Calin-Jageman et al. 2007; Getting 1981). We found that driving one DSI at 20 Hz for either 10 s or 60 s produced a significant increase in the tonic firing rate of other DSIs for ten minutes following the DSI spike train (one way repeated measures ANOVA, F (5, 14) = 15.988, p < 0.001, with post-hoc pairwise comparisons (Holm-Sidak method) vs the pre-DSI spike train firing rate, N = 15 DSIs in 11 preparations; **Figure 3A, B**). While driving a single DSI produced a firing rate elevation for no more than 10 minutes, during an SMP all six DSIs strongly burst for several cycles, which could explain the longer duration (tens of minutes) of the post-SMP elevated DSI firing. Overall, these data support the notion that the DSIs are responsible for their own elevated long-lasting post-SMP tonic firing.

**Figure 3.**
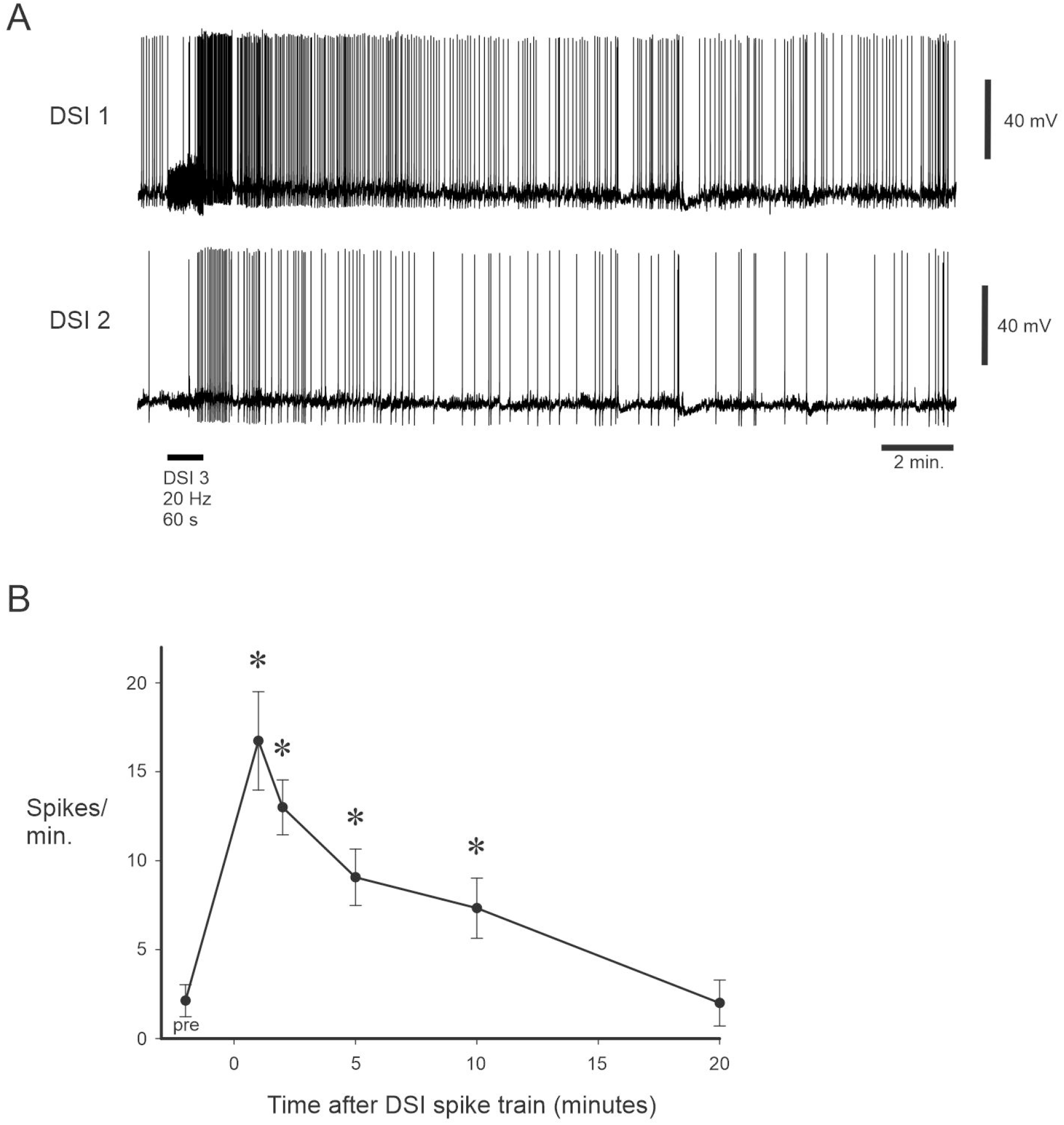
Driving a single DSI causes other DSIs to fire at an elevated rate for several minutes. To further localize the source of the DSI elevated post-SMP firing, we examined the DSIs themselves. **A** Driving a single DSI at 20 Hz for 60 s did not elicit an SMP but did cause an increase in the tonic firing rate of two other DSIs for several minutes. **B** Driving one DSI at 20 Hz for 10 s or 60 s caused other DSIs to significantly increase their firing rate for 10 minutes. This finding reveals that the DSIs are responsible for their own post-SMP elevated tonic firing.

What underlies the ability of the DSIs to induce prolonged tonic firing in one another? For example, are they more excitable after the SMP? To address this question we injected current pulses (0.8 nA for 5 s) three times roughly five seconds apart into DSIs before and at regular intervals (1, 2, 5, 10, 20, 30, 40, 50 and 60 minutes) after an SMP. **Figure 4A** shows an example of DSI excitability before and at 1 and 2 minutes following the SMP. Group data showed that the injected current pulses evoked significantly more spikes in DSIs for 50 minutes post-SMP (one way repeated measures ANOVA, F (9, 8) = 26.34, p < 0.001, with post-hoc pairwise comparisons (Holm-Sidak method) vs pre-SMP, N = 9 DSIs in 5 animals; **Figure 4A, B)**. This increase in DSI excitability correlates well with our prior finding that the DSIs fire at an elevated rate for 55 minutes post-SMP (Popescu and Frost 2002).

**Figure 4.**
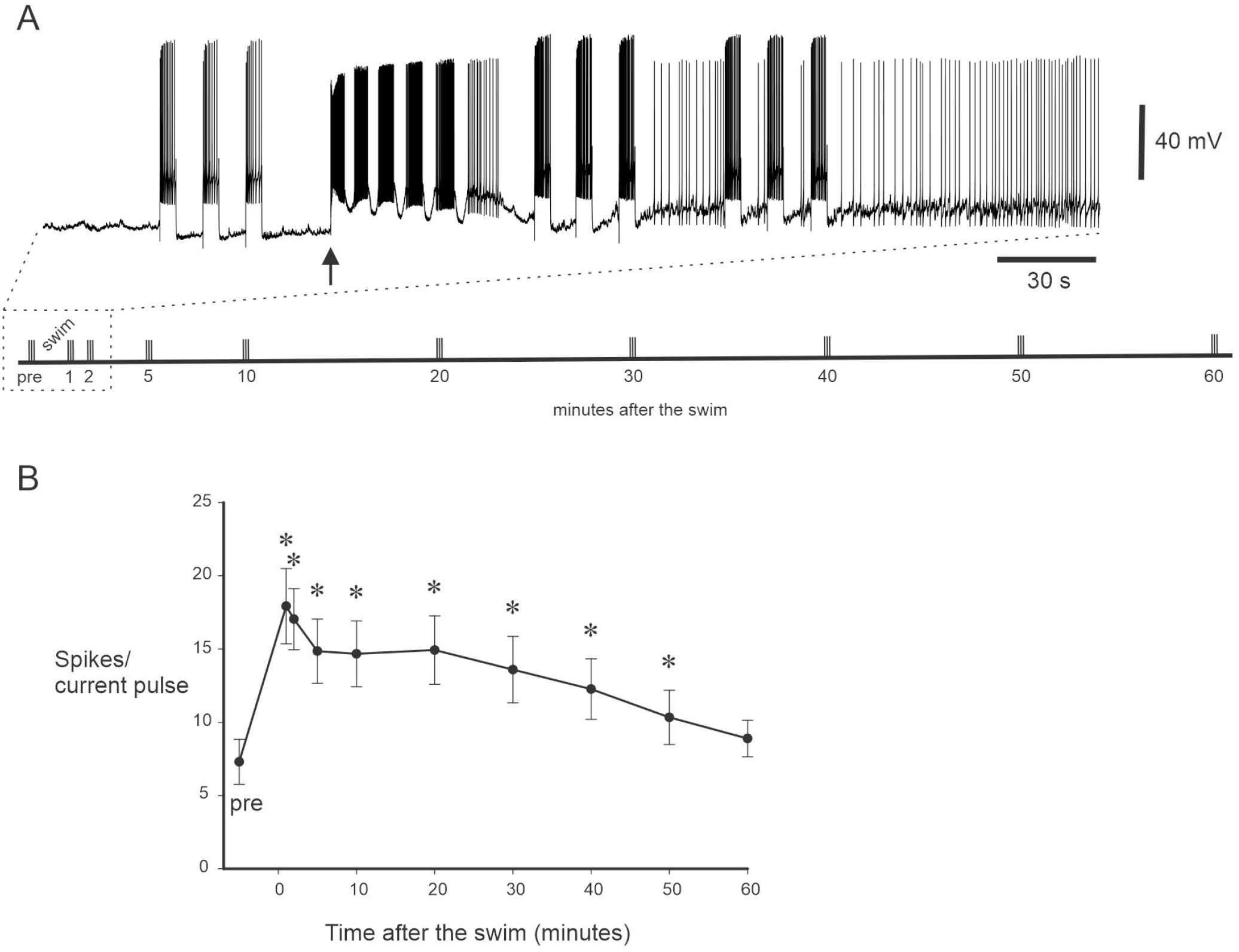
DSI excitability is increased following the SMP for tens of minutes. **A** To examine DSI excitability 0.8 nA, 5 s current pulses were injected into DSIs three times roughly 5 s apart both before and at time points (1, 2, 5, 10, 20, 30, 40, 50 and 60 minutes) following the SMP. The arrow denotes the stimulus to PdN 3 (10V, 10 Hz, 2 s). Inset – schematic of the current injection protocol; the dotted box shows the portion of the protocol shown. **B** DSI excitability was significantly increased for 50 minutes following the SMP.

We then examined whether driving a single DSI could make other DSIs more excitable. As above, three intracellular current pulses (0.8 nA, 5 s) roughly 5 s apart were injected into a single DSI to determine its excitability before and after a one minute 20 Hz spike train in another DSI. **Figure 5A** shows an example of DSI excitability before and at 1 and 2 minutes following such stimulation of another DSI. Note that the DSI 1 spike train resulted in a noticeable membrane potential depolarization in DSI 2. Group data showed that DSIs had significantly elevated excitability for 5 minutes following the spike train in another DSI (one way repeated measures ANOVA, F (9, 4) = 12.14, p < 0.001, with post-hoc pairwise comparisons (Holm-Sidak method) vs pre-DSI spike train, N = 5 DSIs in 4 animals); **Figure 5B)**. These data demonstrate that stimulation of just a single DSI can produce a long-lasting increase in the excitability of other DSIs.

**Figure 5.**
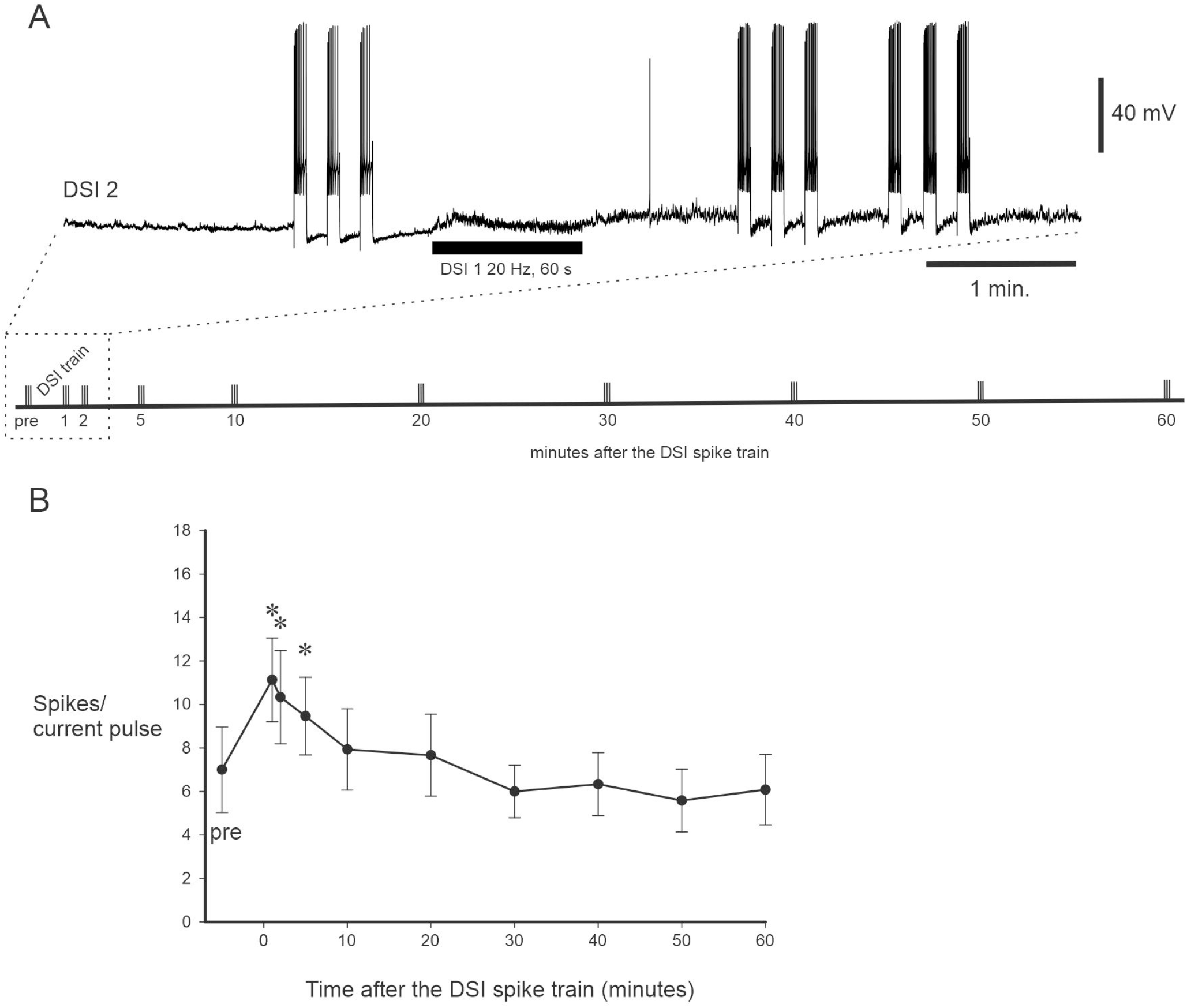
The DSIs enhance each others’ excitability. **A** To examine DSI excitability 0.8 nA, 5 s current pulses were injected into DSIs three times roughly 5 s apart both before and at several time points (1, 2, 5, 10, 20, 30, 40, 50 and 60 minutes) following driving another DSI at 20 Hz for 60 s. Note that the spike train in DSI 1 caused a clear depolarization in DSI 2. Inset – schematic of the current injection protocol; the dotted box shows the portion of the protocol shown. **B** DSI excitability was significantly increased for 5 minutes following the spike train in the other DSI. Thus, even in the absence of an SMP, the DSIs can make each other more excitable for several minutes.

Since neuronal excitability is sensitive to membrane potential, we next examined whether the DSIs are depolarized during the many minutes that they fire at a higher tonic rate following the SMP. Intracellular data were low-pass filtered at 0.1 Hz to reveal the DSI membrane potential with action potentials filtered out. Following SMPs the DSIs were significantly depolarized for 50 minutes (Friedman repeated measures ANOVA on Ranks, Chi-square = 51.74, p =< 0.001, with post-hoc pairwise comparisons (Student-Newman-Keuls), N = 9 DSIs in 6 animals; **Figure 6)**, with the peak depolarization of 10 mV occurring 5 minutes after the SMP. We next tested the direct effect of resting membrane potential on DSI excitability and found that depolarizing DSIs by 10 mV significantly increased their excitability, and conversely, hyperpolarizing them by 10 mV decreased their excitability (One way repeated measures ANOVA, F (2, 4) = 23.07, p < 0.001, with post-hoc pairwise comparisons (Holm-Sidak method) vs control excitability, N = 5 DSIs in 4 animals; control: 8.64±2.74 spikes/current pulse; depolarized by 10 mV: 10.5±2.99 spikes/current pulse; hyperpolarized by 10 mV: 5.5±2.16 spikes/current pulse, (mean±SEM for all)). These data indicate that the post-SMP depolarization of the DSIs likely underlies their increased excitability during the escape crawling period.

**Figure 6.**
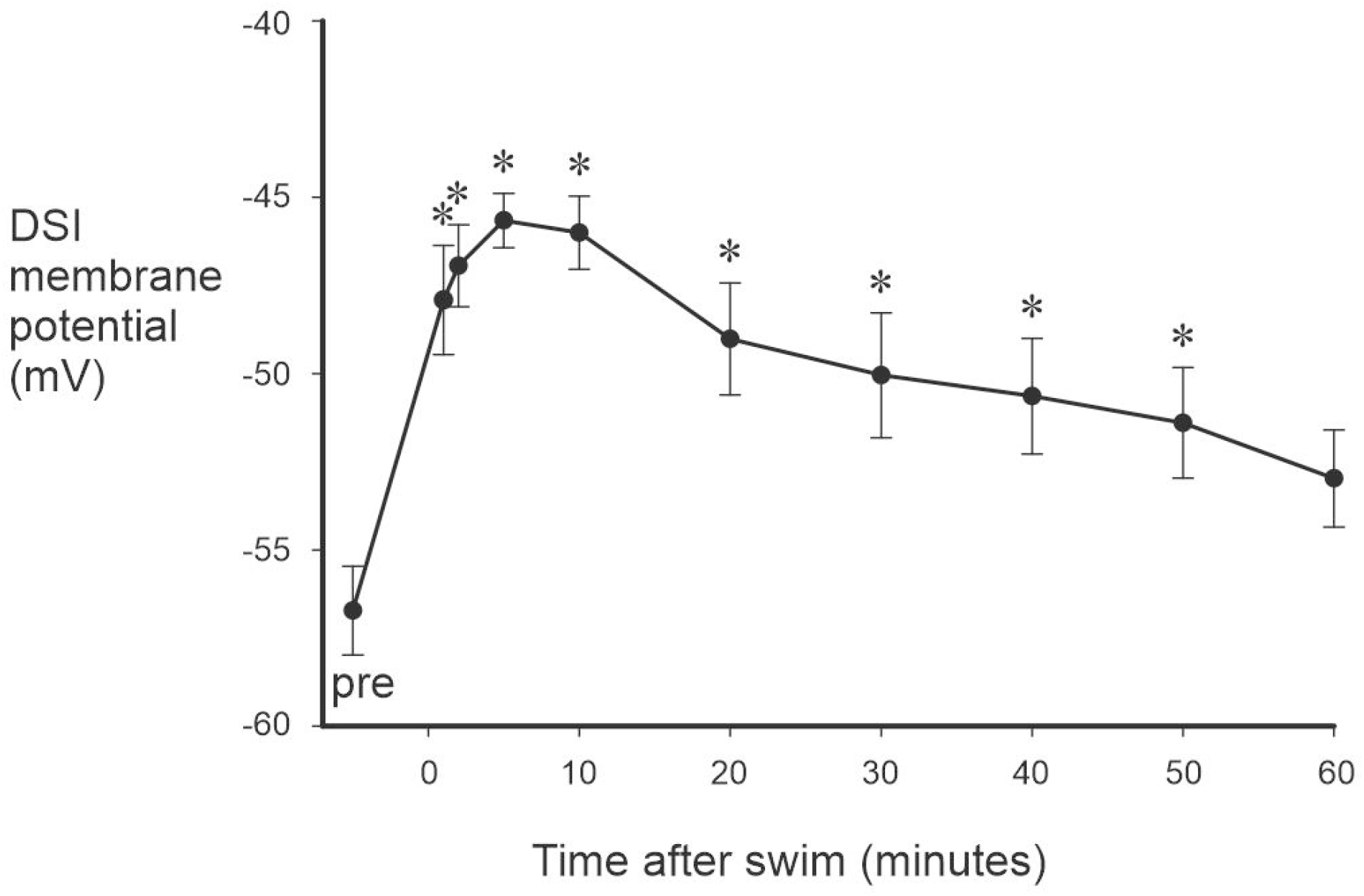
Following an SMP the DSIs are significantly depolarized for 50 minutes. Intracellular data was filtered with a 0.1 Hz low pass filter to remove the spikes to more accurately measure DSI membrane potential. We found that DSI membrane potential was significantly elevated following the SMP for 50 minutes.

### Are the DSIs directly responsible for their elevated post-swim firing?

The DSIs are known to monosynaptically excite one another (Calin-Jageman et al. 2007; Getting 1981), and also to make separable mediating and modulatory excitatory connections onto CPG neuron C2 (Katz and Frost 1995a). This raised the possibility that the DSIs directly modulate one another to produce the prolonged elevated post-SMP tonic firing underlying escape crawling, however this seemed unlikely given the fact that the DSIs polysynaptically inhibit one another in normal saline (Getting and Dekin 1985). Nonetheless, to directly address this question we perfused on a saline solution with a high concentration of divalent cations (Hi Di), which raises the spike threshold and thus minimizes polysynaptic connections (Katz and Frost 1995b). Since Hi Di raises the spike threshold, if the effect of one DSI on another were direct we might not expect to see any increase in DSI spiking following a spike train in another DSI in Hi Di but we would expect to see a membrane potential depolarization caused by the DSI spike train, as seen in **Figure 5A**. A lack of such an effect in Hi Di would be consistent with the effect being indirect, mediated by unknown neurons. In order to visualize membrane potential excursions caused by a DSI spike train, intracellular data were low pass filtered at 0.1 Hz. **Figure 7A** shows that, in control conditions in fASW, a spike train in DSI 1 (20 Hz, 10 s) caused DSI 2 to become depolarized for several minutes. Additionally, the polysynaptic fast inhibition recruited by the DSI 1 spike train is evident in DSI 2 during the spike train. Following Hi Di saline perfusion for 30 minutes, the depolarizing effect of the DSI 1 spike train was completely abolished (**Figure 7B**, N = 4 DSIs in 3 animals), while the polysynaptic inhibition remained. Following washing off of the Hi Di saline for 90 minutes (**Figure 7C**) the depolarizing effect somewhat returned, and the polysynaptic inhibition is present as well. Overall, these data are evidence that the DSIs depolarize each other polysynaptically through as-yet unidentified interneurons.

**Figure 7.**
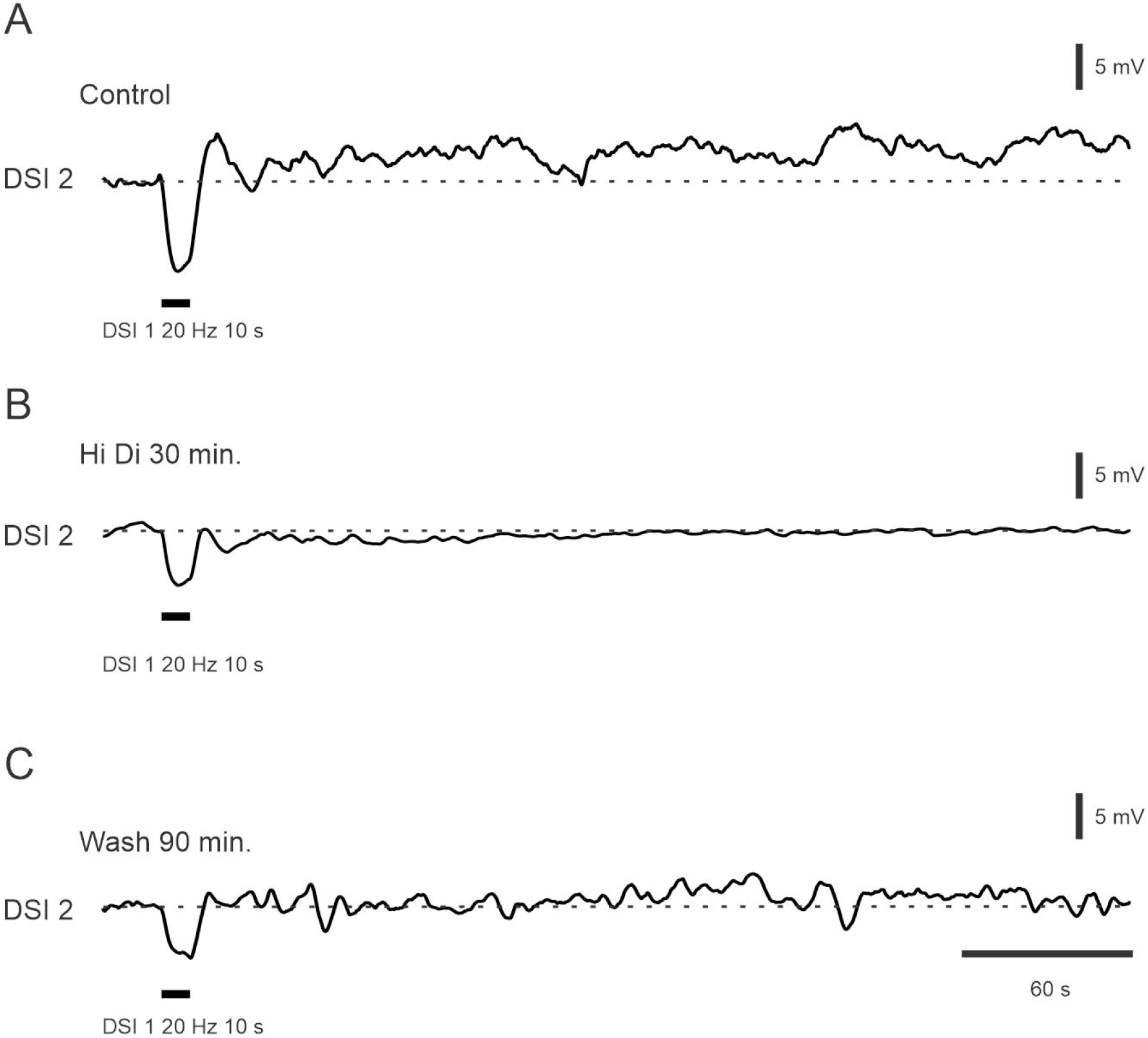
The DSIs’ ability to make each fire at an elevated rate for many minutes is not direct. **A** In fASW driving DSI 1 at 20 Hz for 10 s (trace not shown) causes DSI 2 to become depolarized for several minutes. **B** After perfusion with Hi Di saline for 30 minutes this effect is completely abolished. **C** The ability of the DSI 1 spike train to elicit membrane potential depolarization in DSI 2 somewhat returned following 90 minutes of wash with fASW. Intracellular data was filtered with a 0.1 Hz low pass filter to more clearly visualize DSI membrane potential.

## DISCUSSION

The present study demonstrates a novel network mechanism underlying behavioral sequencing. Since the ability to generate well-coordinated behavioral sequences is essential for survival, their underlying neural mechanisms are of great interest. As elaborated below, several network chain sequencing schemes have been described to date. The most significant contribution of the present study is the demonstration of a novel chain sequencing mechanism, in which generation of the first behavior drives the second. In *Tritonia*, the strong, rhythmic firing of the DSIs that helps to generate the swim behavior also causes a long-lasting depolarization of themselves, which (once the swim is over) makes them more excitable and drives their continuing elevated tonic firing for tens of minutes that excites pedal neurons 5 and 21, two known *Tritonia* crawling neurons (**Figure 1B**). Thus, the crux of the chain sequencing mechanism is that the activity of a specific group of neurons crucial to the generation of the first behavior, the escape swim, induces long-lasting depolarization in themselves, essentially “priming the pump”, to then drive the second behavior, the escape crawl. Unlike prior chain models, rather than relying on propagation through discrete sequentially activated populations with each module going silent once the next is triggered, the swim-to-crawl transition in *Tritonia* appears to arise from activity-dependent persistence within a shared neuromodulatory substrate (**Figure 8**).

**Figure 8.**
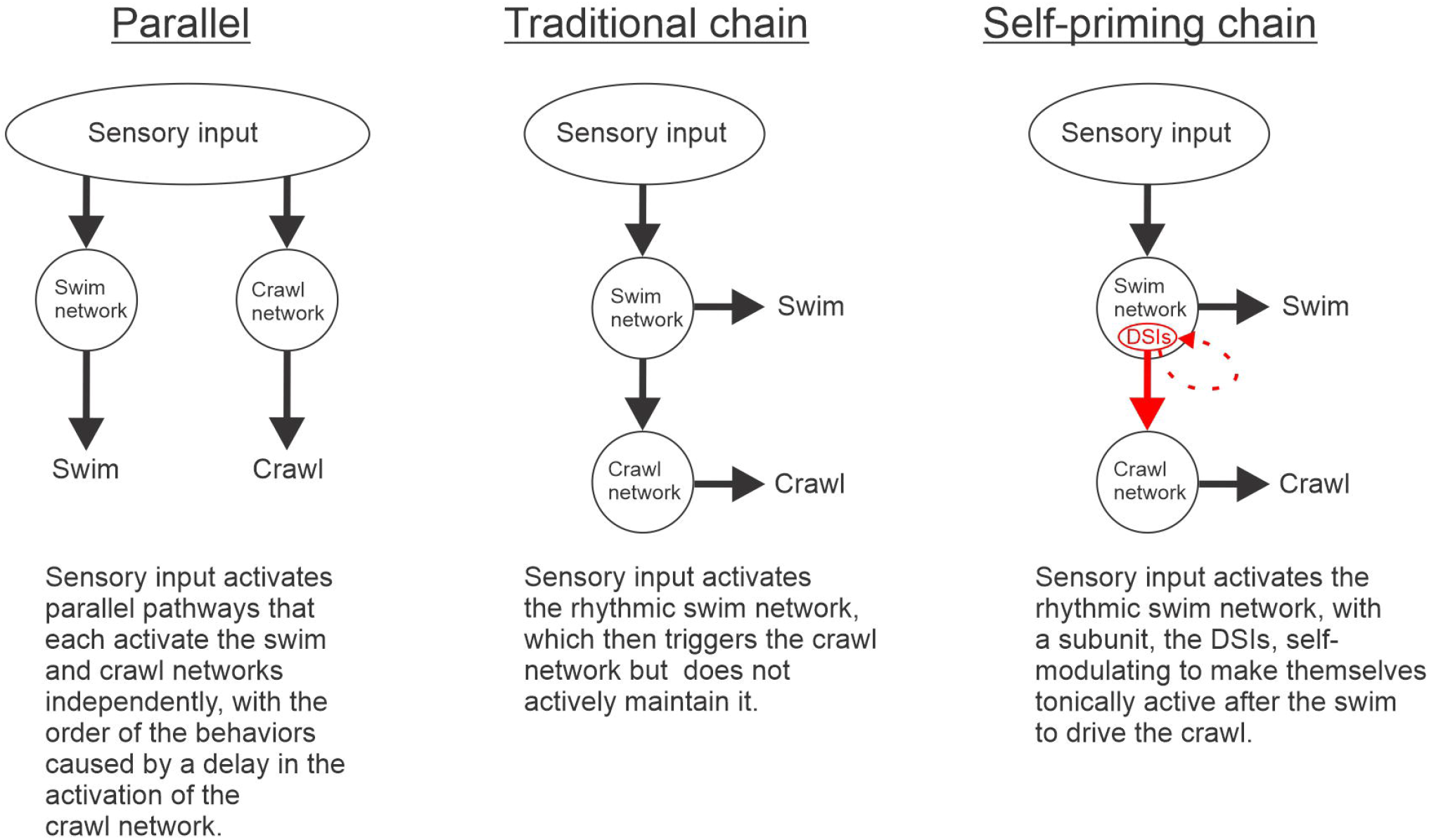
In contrast to parallel or traditional neural chain mechanisms for behavioral sequencing, the *Tritonia* escape swim and crawl behaviors are generated by a novel self-priming neural chain. In a parallel scheme (left) sensory input would activate both the swim and crawl networks in parallel with the sequencing produced by a delay in the onset of the crawl network. In a traditional neural chain (middle), the swim network would trigger the crawl network, but does not actively drive it. In the novel self-priming neural chain mechanism contributed here (right), a module of the swim network (the DSIs) primes itself during the swim (shown as a dashed red line) to then actively drive the escape crawl network for tens of minutes, long after the swim is over.

Currently, the identity of the neurons that mediate the ability of the DSIs to make each other fire at an elevated rate for tens of minutes is unknown. The Hi Di saline results suggest that the DSIs excite as-yet unidentified interneurons that make recurrent excitatory connections back onto the DSIs. Despite the fact that the DSIs act indirectly on one another, they are ultimately responsible for their long-lasting post-swim depolarization that drives the escape crawl.

In recent years, sequential behaviors in several species have been shown to be generated by neural chain sequences. First, a chain sequence involving excitatory and inhibitory neurons has been identified as underlying larval *Drosophila* crawling (Fushiki et al. 2016), which is essentially a peristaltic sequential movement of body segments to move the animal either forwards or backwards. Next, evidence has demonstrated that the larval zebrafish C-turn and escape swim behaviors, two distinct escape behaviors that are always performed in a sequence, are produced by a neural chain involving three different brain regions (Xu et al. 2021). Finally, a feedforward neural chain has been found to underlie sequencing of the syllables of the zebra finch song (Long et al. 2010). Further work identified a role for slow axonal conduction between song bird HVC neurons in the detailed timing of the syllable sequence (Egger et al. 2020). Our present findings represent a novel neural chain mechanism in which generation of the first behavior self-primes the system to generate the second behavior in the sequence.

Additionally, non-chain, parallel activation schemes for the production of sequential behaviors have also recently received experiment support. First, “Ramp-to-threshold” is a type of parallel activation scheme in which successive behaviors are released as different firing thresholds of downstream neurons are reached. A ramp-to-threshold scheme has been shown to underlie various stages of *Drosophila* mating: as the cumulative spike count of a pair of descending neurons, the aSP22 neurons, reaches different thresholds, successive mating behaviors (first proboscis extension, then abdomen bending and finally foreleg lifting) are released (McKellar et al. 2019). Next, a “Parallel activation with hierarchical inhibition” scheme has been found in *Drosophila* sequential grooming – different behavioral modules are simultaneously activated with the order that the behaviors are performed in depending on the order in which inhibition to those modules is lifted (Seeds et al. 2014).

*Tritonia* has been suggested to have evolved from a non-swimming ancestor (Newcomb et al. 2012), with the swimming network having arisen from elements of a pre-existing crawling network. From this perspective it makes sense that the DSIs play dual roles in driving both the escape swimming and crawling behaviors. While previous work showed that the DSIs drive escape crawling via their long-lasting post-SMP elevated tonic firing (Popescu and Frost 2002), our current findings give a mechanistic basis for this enduring post-SMP DSI firing that drives the escape crawl.

Neurons that participate in more than one network/behavior are increasingly being found in many systems. The homolog of C2 in *Pleurobranchaea californica*, the A1 cell, is known to suppress feeding in addition to being a member of the swim CPG (Jing and Gillette 1995). It is therefore possible that C2 may play a similar role in *Tritonia*, and it would be worth exploring whether A1 also drives *Pleurobranchaea* escape crawling. Next, a large number of *Aplysia* pedal ganglion neurons burst rhythmically during both the escape gallop and crawl motor programs (Hill et al. 2020), and *Aplysia* feeding motor programs are generated by a multifunctional network in the buccal ganglion, with different network configurations induced by neuromodulators (Cropper et al. 2023). Further, multifunctional cells are also found in other invertebrate systems such as leeches and crabs – 93% of medicinal leech neurons that oscillate with the swim rhythm also oscillate with the crawl rhythm (Briggman and Kristan 2006), and the crab gastric and pyloric rhythms are generated by a multifunctional stomatogastric ganglion network (Nusbaum et al. 2001). Moreover, a multifunctional network in the mammalian brainstem underlies respiration as well as swallowing, coughing and vomiting (Bianchi and Gestreau 2009). Finally, a recent study showed that there are actually more multifunctional turtle spinal cord neurons that burst rhythmically during both swimming and scratching than there are behaviorally specialized neurons (Morris et al. 2024).

It is worth noting that the post-SMP elevated tonic firing of the DSIs also serves another purpose: retaining the memory for sensitization of the escape swim (Hill et al. 2015), so that if the animal is attacked again, it will start the escape swim with a shorter latency, it will have a more forceful initial “jump” off of the substrate (Hill et al. 2015; Mistry et al. 2026), and in some instances it can perform more swim cycles (Frost et al. 1998). Thus not only are the DSIs multifunctional in the sense that they drive both escape swimming and crawling, their post-SMP elevated tonic firing does two things at once: it drives escape crawling and simultaneously maintains the memory for sensitization of the escape swim.

In summary, we have provided evidence that a self-priming neural chain involving identified serotonergic CPG neurons drives the sequential *Tritonia* escape swim/crawl behaviors. In particular, prolonged self-induced post-SMP depolarization of the DSIs is the cellular mechanism driving the second behavior of the sequence. In contrast to parallel schemes and traditional feedforward neural chains, our data have revealed a novel type of neural chain in which a subunit of the first network modulates itself to produce long-lasting drive to the neurons generating the second behavior in the sequence (**Figure 8**).

We have found that the same neurons (the DSIs) participate continuously across both behaviors while their activity-dependent change in state links the two actions into a coordinated sequence. Our finding represents a simple and elegant mechanism by which the nervous system generates sequential actions across timescales.

## REFERENCES

Audesirk G. Central neuronal control of cilia in Tritonia diamedia. Nature 272: 541–543, 1978.

Bianchi AL, and Gestreau C. The brainstem respiratory network: an overview of a half century of research. Respir Physiol Neurobiol 168: 4–12, 2009.

Briggman KL, and Kristan WB, Jr. Imaging dedicated and multifunctional neural circuits generating distinct behaviors. J Neurosci 26: 10925–10933, 2006.

Calin-Jageman RJ, Tunstall MJ, Mensh BD, Katz PS, and Frost WN. Parameter space analysis suggests multi-site plasticity contributes to motor pattern initiation in Tritonia. J Neurophysiol 98: 2382–2398, 2007.

Clower WT. Early contributions to the reflex chain hypothesis. J Hist Neurosci 7: 32–42, 1998.

Cropper EC, Perkins M, and Jing J. Persistent modulatory actions and task switching in the feeding network of Aplysia. Curr Opin Neurobiol 82: 102775, 2023.

Delcomyn F. Neural basis of rhythmic behavior in animals. Science 210: 492–498, 1980.

Egger R, Tupikov Y, Elmaleh M, Katlowitz KA, Benezra SE, Picardo MA, Moll F, Kornfeld J, Jin DZ, and Long MA. Local Axonal Conduction Shapes the Spatiotemporal Properties of Neural Sequences. Cell 183: 537–548 e512, 2020.

Frost WN, Brandon CL, and Mongeluzi DL. Sensitization of the Tritonia escape swim. Neurobiol Learn Mem 69: 126–135, 1998.

Frost WN, and Katz PS. Single neuron control over a complex motor program. Proc Natl Acad Sci U S A 93: 422–426, 1996.

Fushiki A, Zwart MF, Kohsaka H, Fetter RD, Cardona A, and Nose A. A circuit mechanism for the propagation of waves of muscle contraction in Drosophila. Elife 5: 2016.

Getting PA. Mechanisms of pattern generation underlying swimming in Tritonia. I. Neuronal network formed by monosynaptic connections. J Neurophysiol 46: 65–79, 1981.

Getting PA. Mechanisms of pattern generation underlying swimming in Tritonia. III. Intrinsic and synaptic mechanisms for delayed excitation. J Neurophysiol 49: 1036–1050, 1983a.

Getting PA. Neural control of swimming in Tritonia. Symp Soc Exp Biol 37: 89–128, 1983b.

Getting PA, and Dekin MS. Mechanisms of pattern generation underlying swimming in Tritonia. IV. Gating of central pattern generator. J Neurophysiol 53: 466–480, 1985.

Hill ES, Brown JW, and Frost WN. Photodiode-Based Optical Imaging for Recording Network Dynamics with Single-Neuron Resolution in Non-Transgenic Invertebrates. J Vis Exp 2020.

Hill ES, Vasireddi SK, Wang J, Bruno AM, and Frost WN. Memory Formation in Tritonia via Recruitment of Variably Committed Neurons. Curr Biol 25: 2879–2888, 2015.

Hill ES, Wang J, Brown JW, Mistry VK, and Frost WN. Surprising multifunctionality of a Tritonia swim CPG neuron: C2 drives the early phase of postswim crawling despite being silent during the behavior. J Neurophysiol 132: 96–107, 2024.

Jing J, and Gillette R. Neuronal elements that mediate escape swimming and suppress feeding behavior in the predatory sea slug Pleurobranchaea. J Neurophysiol 74: 1900–1910, 1995.

Katz PS, and Frost WN. Intrinsic neuromodulation in the Tritonia swim CPG: serotonin mediates both neuromodulation and neurotransmission by the dorsal swim interneurons. J Neurophysiol 74: 2281–2294, 1995a.

Katz PS, and Frost WN. Intrinsic neuromodulation in the Tritonia swim CPG: the serotonergic dorsal swim interneurons act presynaptically to enhance transmitter release from interneuron C2. J Neurosci 15: 6035–6045, 1995b.

Long MA, Jin DZ, and Fee MS. Support for a synaptic chain model of neuronal sequence generation. Nature 468: 394–399, 2010.

McKellar CE, Lillvis JL, Bath DE, Fitzgerald JE, Cannon JG, Simpson JH, and Dickson BJ. Threshold-Based Ordering of Sequential Actions during Drosophila Courtship. Curr Biol 29: 426–434 e426, 2019.

Mistry VK, Hill ES, and Frost WN. Strategies used by two memories to share space in a small molluscan neural network. iScience 29: 115355, 2026.

Morris MM, Hao 郝赵哲 ZZ, and Berkowitz A. Electrophysiological Activity of Multifunctional and Behaviorally Specialized Spinal Neurons Involved in Swimming, Scratching, and Flexion Reflex in Turtles. eNeuro 11: 2024.

Newcomb JM, Sakurai A, Lillvis JL, Gunaratne CA, and Katz PS. Homology and homoplasy of swimming behaviors and neural circuits in the Nudipleura (Mollusca, Gastropoda, Opisthobranchia). Proc Natl Acad Sci U S A 109 Suppl 1: 10669–10676, 2012.

Nusbaum MP, Blitz DM, Swensen AM, Wood D, and Marder E. The roles of co-transmission in neural network modulation. Trends Neurosci 24: 146–154, 2001.

Popescu IR, and Frost WN. Highly dissimilar behaviors mediated by a multifunctional network in the marine mollusk Tritonia diomedea. J Neurosci 22: 1985–1993, 2002.

Seeds AM, Ravbar P, Chung P, Hampel S, Midgley FM, Jr., Mensh BD, and Simpson JH. A suppression hierarchy among competing motor programs drives sequential grooming in Drosophila. Elife 3: e02951, 2014.

Sherrington C. The integrative action of the nervous system. New Haven, CT: Yale University Press, 1947.

Willows AO. Physiological basis of feeding behavior in Tritonia diomedea. II. Neuronal mechanisms. J Neurophysiol 44: 849–861, 1980.

Xu L, Guan NN, Huang CX, Hua Y, and Song J. A neuronal circuit that generates the temporal motor sequence for the defensive response in zebrafish larvae. Curr Biol 31: 3343–3357 e3344, 2021.

